# Calibrating developmental dynamics from single-cell atlases: lineage paths outrun least-action routes

**DOI:** 10.64898/2026.11.05.764549

**Authors:** Qin Liu, Ying Zhou, Jielin Cao, Yiqiao Wang

## Abstract

Single-cell atlases infer developmental dynamics from destructive snapshots, yet how far realized lineage paths depart from such inferred dynamics has never been measured with controls. We introduce LADD (Least-Action Developmental Dynamics), a measurement framework that learns a stochastic developmental vector field from time-series single-cell RNA-seq and computes minimum-action and minimum-time paths, together with a three-layer calibration protocol that separates genuine path deviation from the generic cost of centroid measurement, and we compare its forecasting component with optimal-transport and flow-matching methods. In the lineage-resolved *Caenorhabditis elegans* atlas, the raw action ratio of observed lineage-centroid trajectories to optimal paths is 3.79, yet forward-simulated ensembles of the learned model itself score 2.07, leaving a genuine excess of about two-fold. In zebrafish (*Danio rerio*), the observed ratio sits entirely at the model’s floor. Three mechanisms that could explain the residual deviation (detours, flow-opposition, and fate-map complexity) are each excluded; what remains is speed, with lineages moving 30–43% faster along the learned flow. We report validity boundaries for temporal resolution, cross-species comparison, and energetic interpretation, delimiting where action-based trajectory comparisons are meaningful.

**Author Summary:** Embryos develop from a single cell into many specialized cell types, and modern atlases let researchers watch this process as a movie of molecular snapshots. Because each cell is measured only once, software must stitch snapshots into “trajectories”, and some studies ask whether development follows optimal routes, such as paths of least action (a principle borrowed from physics). Here we provide LADD, a method that learns the dynamics of development from such atlases and computes the optimal paths those dynamics imply, together with a calibration protocol that distinguishes genuine biological deviation from artifacts created by measuring averages of cell populations. Applying it to worm and zebrafish embryogenesis, we find that most of the apparent deviation from optimality is such a measurement artifact; what remains in the worm is a modest, genuine excess arising because lineages move faster than the optimal route, not around it. LADD offers a reusable way to test optimality claims about development against an explicit noise floor.

## 1. Introduction

Single-cell RNA-seq time courses now let us watch embryogenesis as a movie of cell states, and dynamics-inference methods (RNA velocity [1], optimal transport [2], flow matching [3,4], and successors) turn snapshot movies into vector fields, landscapes, and predicted trajectories [5,6]. Every reconstruction rests on destructive sampling: each cell is measured once, so a “trajectory” is a population summary stitched across embryos, not the real path of any cell. How far, then, do realized developmental paths depart from the routes that inferred dynamics consider natural? Despite the field’s reliance on these constructions, the departure has never been measured with controls.

The sharpest version of this question comes from physics. Since Bernoulli’s 1696 brachistochrone problem, variational principles (least action, least time) have described nature’s paths as optima, and Waddington’s epigenetic landscape [7] invites the same reading for development. Recent work has turned the metaphor into explicit claims: embryogenesis as variational free-energy minimization [8], growing tissues minimizing spatiotemporal growth variation [9], development assumed to follow minimum-action paths [10]. These proposals *assume* an optimality principle rather than test it: none computes the optimal path through a learned developmental dynamics and compares it, with controls, to the paths embryos actually take.

A fastest-development law also faces a basic objection. Physical variational principles minimize a single hard objective, but developmental timing is shaped by selection on fitness, of which speed is only one component: transition timings are optimized against growth and mortality [11,12], developmental time scales with body mass and temperature [13], and diapause delays development adaptively. The path is equally debatable: canalization shapes valleys for robustness, not speed, and morphogenesis is an active, far-from-equilibrium process, though some modules are provably time-optimal [14] and early cell cycles trade checkpoint stringency for speed [15,16]. Judging such claims, a decades-old debate [17], requires an instrument that measures how close a realized trajectory comes to the optimum and distinguishes “shaped by selection” from “truly optimal” [18].

Here we introduce **LADD** (Least-Action Developmental Dynamics), which couples a neural stochastic differential equation learned from time-series single-cell RNA sequencing (scRNA-seq) with Freidlin–Wentzell large-deviation theory [19] to compute minimum-action and minimum-time developmental paths. We apply it to the lineage-resolved *C. elegans* embryogenesis atlas [20], whose invariant, fully mapped cell lineage provides a unique ground truth, and we replicate the core measurements on the zebrafish embryogenesis atlas [21]. The central lesson is methodological as much as biological: because single-cell profiling is destructive, developmental trajectories are measured as the paths of population *centroids*, and centroid paths carry a generic, calibratable excess action unrelated to the route any cell actually takes. Most of the raw deviation from least-action paths is this measurement floor; what remains after calibration is a ≈2-fold path-specific excess in the worm, generated by lineages moving *faster* than the least-action route rather than around it. We report the calibration protocol, a comparison of the learned dynamics with optimal-transport and flow-matching methods, the calibrated measurement, and the validity boundaries (temporal resolution, cross-species comparability, energetic interpretation) within which action-based trajectory comparisons are meaningful. LADD is released as a reusable package, with trained checkpoints and per-path source data sufficient to reproduce every figure.

## 2. Results

We first define the measurement framework and its calibration protocol, then validate the instrument against held-out data and null models, and finally apply the calibrated instrument to characterize the excess action in *C. elegans*.

### 2.1 The LADD framework: a learned stochastic dynamical model of embryogenesis

We analyzed the lineage-resolved *C. elegans* embryogenesis atlas of Packer et al. [20], retaining 87,702 high-quality cells spanning 150–750 min after first cleavage (Fig 1A–D), embedded in a 50-dimensional principal-component analysis (PCA) space of 3,000 highly variable genes (29.9% variance explained). Batch composition was well mixed in early and mid bins; the latest bins showed moderate enrichment (Fig 1F), a caveat for late-time interpretation.

**Fig 1.**
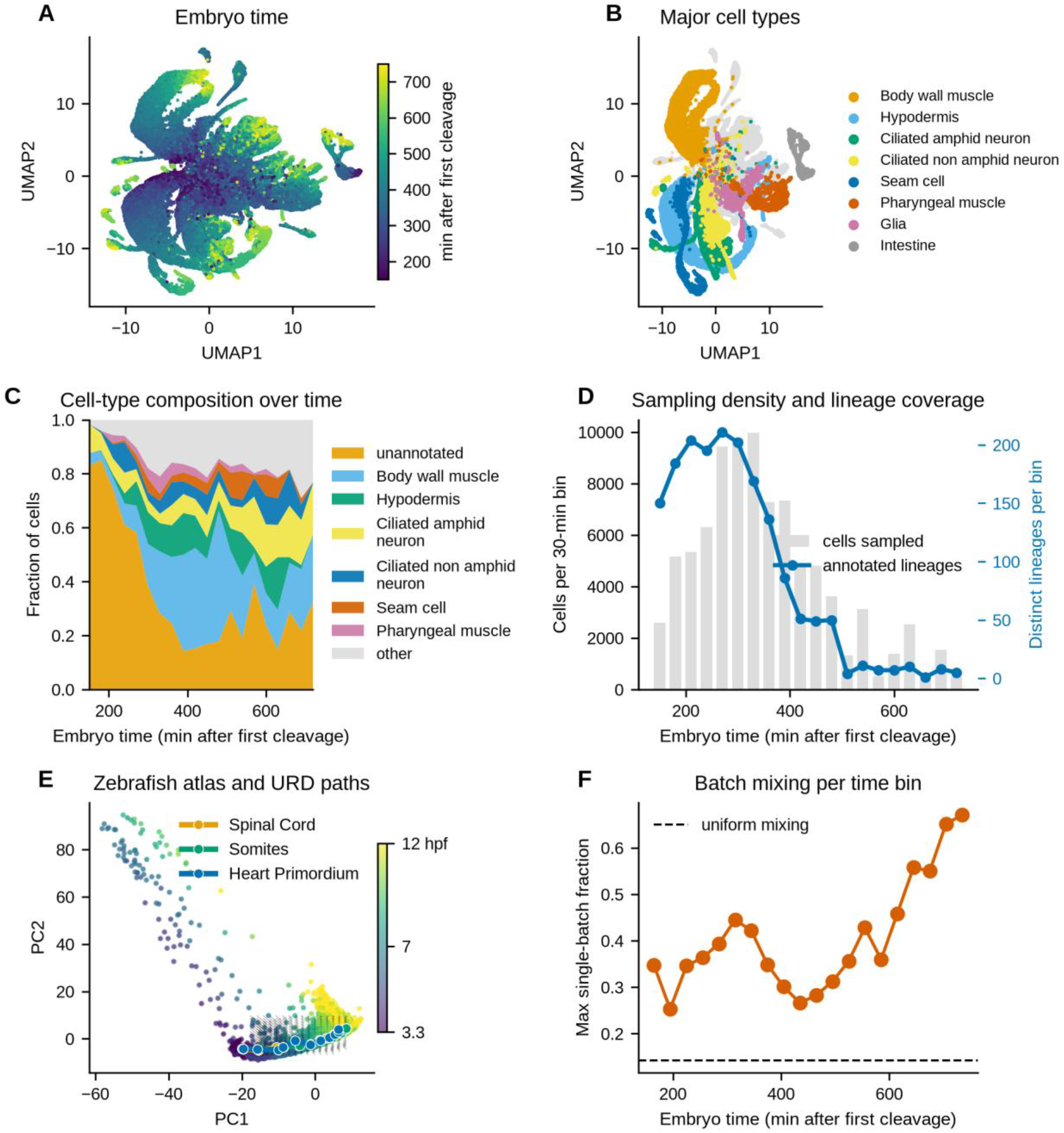
Two lineage-resolved embryogenesis atlases. (A) Uniform manifold approximation and projection (UMAP) of the 87,702 *C. elegans* cells analyzed (150–750 min after first cleavage), colored by embryo time. (B) Same embedding, colored by major cell type (eight most frequent; rest in gray). (C) Cell-type composition over developmental time (stacked fractions per 30-min bin). (D) Sampling density (bars: cells per 30-min bin) and lineage coverage (blue line: distinct annotated lineages per bin); coverage falls after ∼550 min. (E) Zebrafish atlas (38,731 cells, 3.3–12 h post-fertilization) in PCA space, colored by stage, with the learned drift field (arrows) and three example URD branch centroid paths. (F) Batch mixing: maximum single-batch fraction of cells per 30-min bin (dashed line: uniform mixing across seven batches).

LADD learns an autonomous stochastic vector field dx = f(x)dt + Σ^(1/2)dW in this latent space (S1A Fig). The drift f is a neural network trained to match empirical one-step displacements between adjacent 30-min bins via local k-nearest-neighbor (k-NN) coupling (k = 10); the diffusion covariance Σ is calibrated so the one-step-ahead marginal spread matches the data (Methods). Local coupling is essential here: global entropic optimal-transport couplings yield blurred, divergent rollouts (S1 Note). A scalar potential U(x) fitted to the drift (Helmholtz-style decomposition, Methods) yields an interpretable landscape (S1B Fig); the field is predominantly gradient-like (median rotational fraction 17% per cell; S1C,D Fig).

The learned field is predictive and biologically coherent, serving as the reference dynamics below. It beat persistence and linear-drift baselines in 11 of 13 evaluable interior held-out bins and at 60–90-min forward extrapolation (S3 and S4 Figs; S4 Note). Predicted cell futures respect the invariant lineage: sister lineages’ rolled-out futures are 6.3× closer than random pairs (p ≈ 10⁻⁷), and mid-development rollouts improve terminal-fate prediction over persistence (S2 Fig; S5 Note). The same pipeline fitted to the zebrafish atlas of Farrell et al. [21] (38,731 cells, 12 stages, 3.3–12 h post-fertilization; Fig 1E) passed the same checks (S3 Note).

### 2.2 Developmental trajectories deviate substantially from least-action paths

We reconstructed 179 lineage-branch trajectories (founder-to-terminal, median 9 time-bins each) as centroid paths through latent space. For each path, we computed the Freidlin– Wentzell action S = ½∫|ẋ − f(x)|²_Σ⁻¹ dt, which measures how improbable a path is under the learned dynamics; the minimum-action path between two endpoints is the most probable route. We compared each observed trajectory to the minimum-action path between the same endpoints and duration, with controls.

Three findings emerged (Fig 2A; S8A–C Fig). **First**, observed trajectories are far from least-action: the median action ratio S_observed/S_min was ≈3.9 (interquartile range [IQR] 2.7–5.9), and even a straight-line route carried only ≈2× the minimum action (S_observed/S_line ≈ 1.9). **Second**, the excess is not random noise. Shuffling the temporal order of waypoints raised the action ≈4.2-fold, while time reversal raised it only ≈1.2-fold; the realized *sequence* of states is dynamically coherent even though its *route* departs from the optimum. **Third**, the excess is structured: largest for shallow, founder-level branches (median ≈5.0 vs ≈3.4 for deep branches; S8D Fig) and strongly correlated with path curvature (Spearman ρ = 0.60, p = 5.8×10⁻¹⁹).

**Fig 2.**
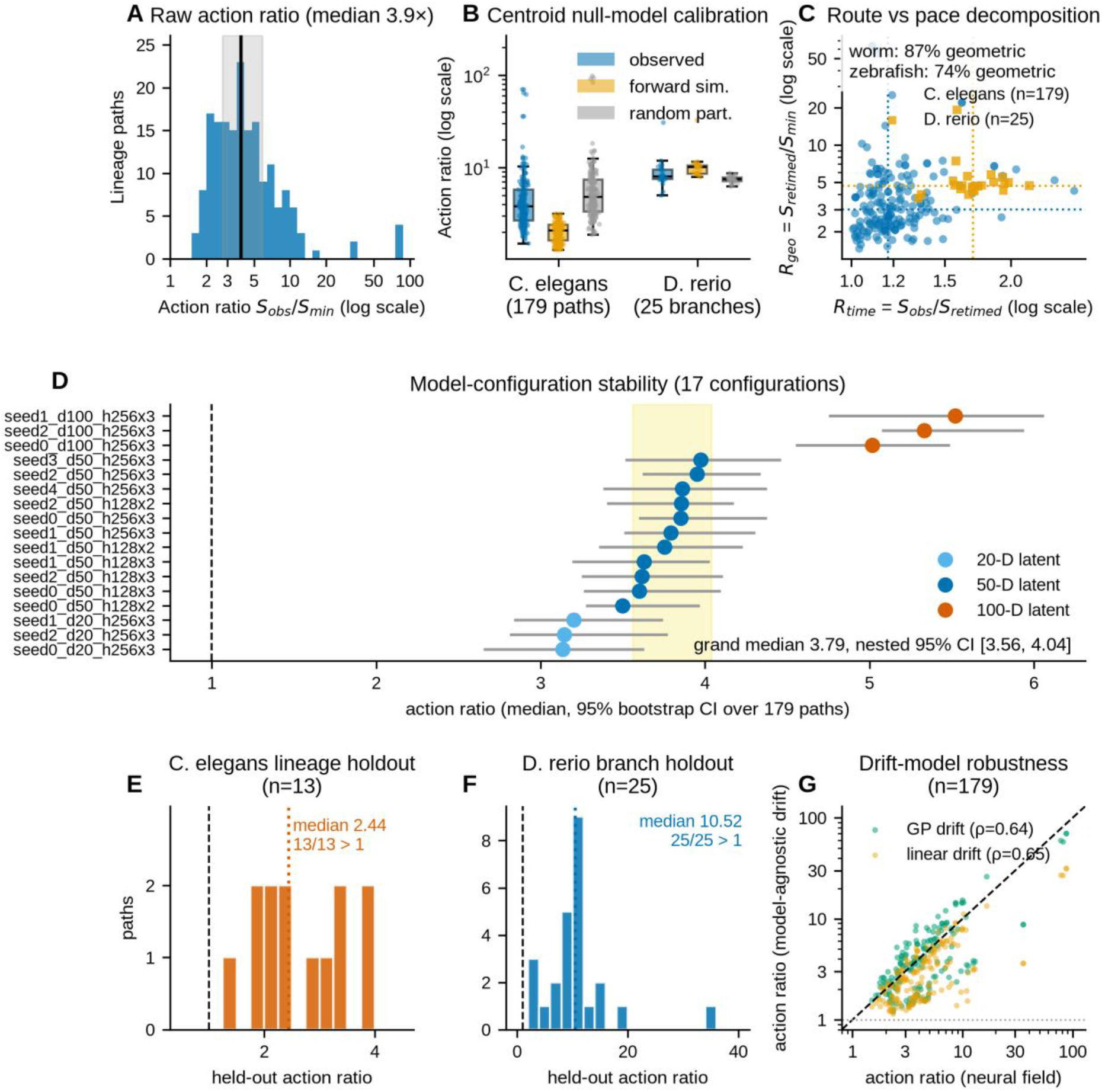
Calibrating the action ratio. (A) Distribution of the action ratio S_observed/S_min across 179 *C. elegans* lineage-branch paths (log scale; line: median 3.9×; band: IQR). (B) Centroid null-model calibration: action ratios of observed centroid paths, forward-simulated ensembles (2,000 particles per branch under the learned field), and random-partition centroids (matched cell counts, lineage structure destroyed), for *C. elegans* (179 paths) and *D. rerio* (25 branches, post-epiboly); log scale. Worm observed paths lie above the simulated floor (median 3.79 vs 2.07, Wilcoxon p = 4×10⁻³¹) but below random partitions (4.77, p = 3×10⁻⁵); zebrafish observed paths sit at the floor (7.95 vs 10.03, p = 5×10⁻³). (C) Geometry-versus-timing decomposition: timing factor R_time = S_observed/S_retimed versus geometry factor R_geo = S_retimed/S_min per path (log–log; dotted lines: medians); 87% (worm) and 74% (zebrafish) of the log-excess is geometric. (D) Median manifold-constrained action ratio (points) with 95% bootstrap CI over paths for 17 model configurations (five reference seeds; 128×2/128×3 networks; 20-D/100-D latent spaces; points colored by dimensionality). Shaded band: grand median 3.79, nested-bootstrap 95% CI [3.56, 4.04]; dashed line: ratio = 1 (least action). (E) *C. elegans* lineage holdout: action ratios of the 13 deep branch paths whose lineages were withheld from field training (median 2.44; 13/13 > 1). (F) *D. rerio* leave-one-branch-out: held-out ratios for all 25 URD branches (median 10.52; 25/25 > 1). (G) Model-agnostic drift surrogates: per-path action ratios under a Gaussian-process drift (green) and a global linear drift (orange) versus the neural field (n = 179; log–log; dashed: identity; dotted: ratio = 1; Spearman ρ = 0.64, 0.65); all paths exceed 1 under all three drifts.

Unconstrained minimization could exploit regions far from the data manifold, but recomputing minimum-action paths under a manifold penalty left the ratio unchanged (median 3.85 constrained vs 3.88 unconstrained; ρ = 0.9998; S8E Fig; S7 Note), so the deviation is not an off-manifold artifact. Minimum-action paths also sit no closer to the observed route than a straight line does (median Chamfer distance 3.5 vs 3.1), but this set-level comparison cannot distinguish a different route from the same route at different pace; Section 2.10 resolves this.

We decomposed the action multiplicatively: for each observed path, we computed the exact optimal re-timing of the same curve under the fixed duration (closed-form Karush–Kuhn–Tucker (KKT) condition; Methods 4.6), giving R_obs = R_time × R_geo, where R_time = S_obs/S_retimed penalizes the observed time allocation and R_geo = S_retimed/S_min measures the residual excess from geometry alone (Fig 2C). In the worm, R_time = 1.17 (IQR 1.10–1.29) while R_geo = 3.01 (IQR 2.29–4.63): 87% of the log-excess is geometric, and no path (0/179) is timing-dominated. Zebrafish post-epiboly paths show the same pattern with a larger timing share (R_time = 1.70, IQR 1.61–1.84; R_geo = 4.68, IQR 4.39–4.99; 74% geometric). The excess is therefore primarily a property of *where* trajectories go, and only secondarily of uneven tempo.

### 2.3 A three-layer calibration protocol: stability, circularity controls, and the ensemble floor

Before interpreting the ratio biologically, it must survive retraining, resist circularity, and be compared against what the same measurement assigns to null ensembles: three calibration layers.

#### Layer 1: statistical and architectural stability

We re-estimated the full pipeline under 17 model configurations spanning training seeds, network size, and latent dimensionality, recomputing all 179 path ratios per configuration (Fig 2D). The grand median ratio is 3.79 (nested-bootstrap 95% CI [3.56, 4.04]); every configuration median exceeds 3.1, and per-path rankings are stable (Spearman ρ = 0.77–0.99 against the reference seed). One sensitivity emerged: the ratio grows monotonically with latent dimensionality (≈3.2 in 20-D, 3.79 in 50-D, 5.0–5.5 in 100-D), because minimum-action paths find shorter routes in higher dimensions. The trend is smooth and qualitative conclusions hold at every dimensionality. The 50-D headline is not tuned to the action: forecasting skill and variance explained improve with dimensionality (relative W2 [2-Wasserstein distance] 0.87/0.83/0.80; 22/30/37%) while rollout lineage coherence degrades (7.9×/6.6×/5.1×); the 50→100-D forecast gain is small (3%) against a 24% coherence loss, and 50-D matches both source atlases (S9A,B Fig).

#### Layer 2: circularity controls

The same data train the field and define the paths, and the field could encode the excess by construction. First, lineage holdout: retraining the field with one-fifth of terminal lineages withheld gave a median ratio of 2.44 for the 13 fully held-out deep branch paths (13/13 > 1), versus 3.88 with full data (Fig 2E); the excess survives, reduced as expected for interpolation into unseen territory. Zebrafish leave-one-branch-out retraining gave a median held-out ratio of 10.52 (25/25 > 1; Fig 2F), against 8.70 with the full model. Second, model-agnostic drift: replacing the neural drift with a global linear drift or a Gaussian-process drift, both far less predictive (R² = 0.15 and 0.11 vs 0.40), still reproduced the excess (medians 2.39 and 3.56; all 179 paths > 1; per-path ρ = 0.65 and 0.64 with the neural ratios; Fig 2G). The excess is therefore parameterization-independent in sign and rank, though its absolute magnitude (2.4– 3.9 across drifts) is field-dependent. A time-conditioned field f(x, t) gave an identical picture (median ratio 3.68 vs 3.86; ρ = 0.94).

#### Layer 3: the ensemble floor

The centroid of a spreading, bifurcating ensemble is not the trajectory of any cell, so the action functional may assign a high ratio to *any* centroid path. We calibrated this floor directly (Fig 2B; Methods 4.15). First, synthetic ground truth: ensembles simulated under four exactly solvable 50-dimensional scenarios (static diffusion, coherent translation, a 60/40 bifurcation, an asynchronous wave), analyzed with the identical pipeline, gave centroid ratios of 1.5–3.0: perfectly coherent dynamics do not score exactly 1. Second, forward simulation: ensembles of 2,000 particles evolved under the learned field from each branch’s start population gave centroid ratios of 2.07 (IQR 1.61–2.39) in worm and 10.0 (IQR 8.54–10.41) in zebrafish. Third, random-partition centroids (matched cell counts, lineage structure destroyed) scored 4.77 (IQR 3.29–7.33) in worm and 7.50 (IQR 7.09–7.74) in zebrafish. Against these references, observed worm paths (3.79) lie significantly *above* the model’s own floor (median per-path fold 1.96; Wilcoxon p = 4×10⁻³¹) and significantly *below* random partitions (p = 3×10⁻⁵). About half of their raw excess is therefore the generic price of being a centroid, not a property of the realized route. In zebrafish, observed ratios sit at or below the simulated floor (7.95 vs 10.03, p = 5×10⁻³; Fig 3E) and are indistinguishable from random partitions, so we do not interpret the zebrafish excess as route suboptimality.

**Fig 3.**
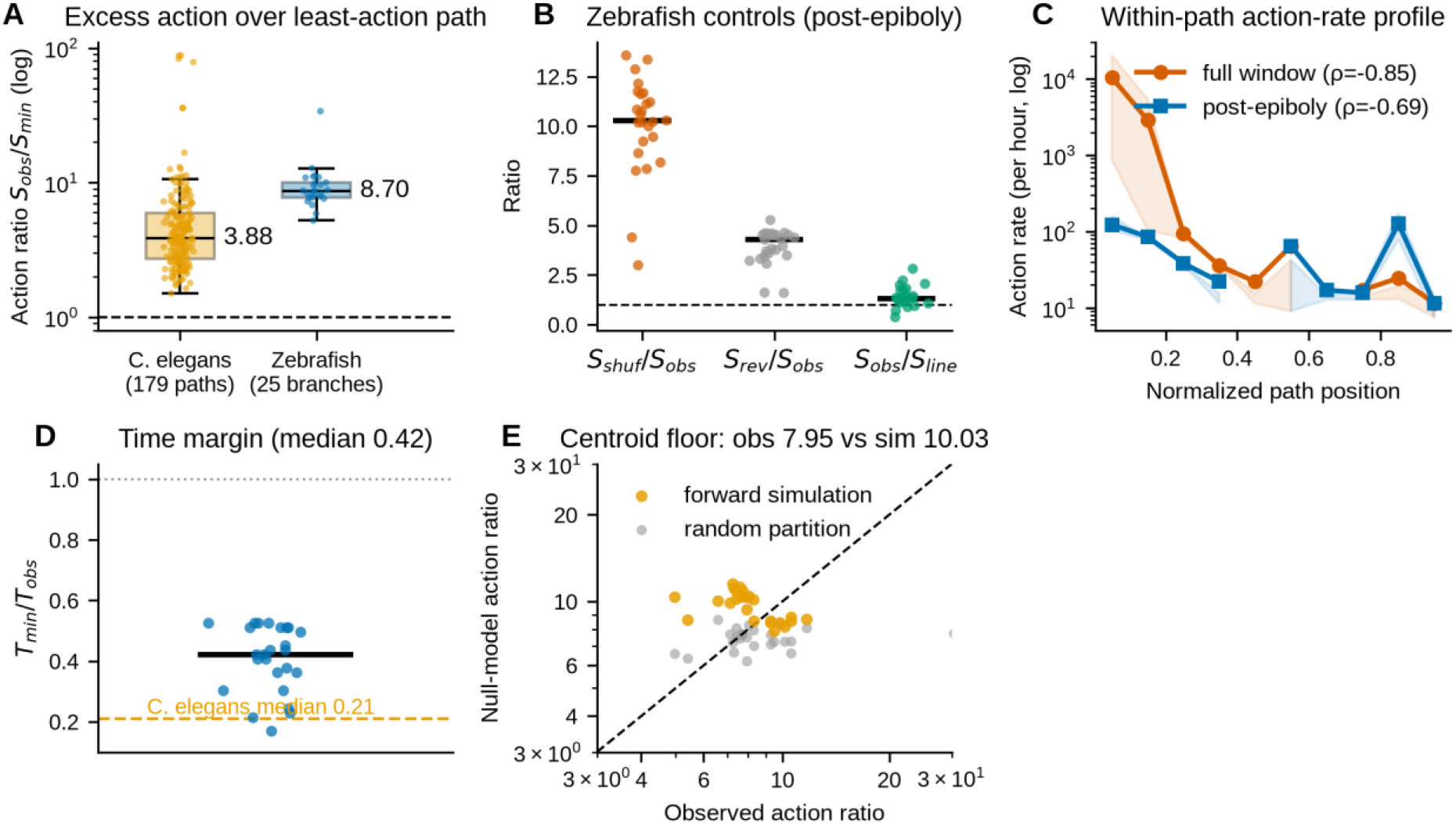
Cross-species comparison in zebrafish embryogenesis. (A) Action ratios S_observed/S_min for 179 *C. elegans* lineage-branch paths (orange) and 25 zebrafish URD branch paths over the post-epiboly window (blue); medians annotated. (B) Control ratios for the 25 zebrafish branches: waypoint-shuffled (red), time-reversed (gray), and straight-line (green) versus observed; black bars: medians. (C) Within-path action-rate profiles: full window (red; dominated by the epiboly burst, Spearman ρ = −0.85) and post-epiboly window (blue); binned medians with interquartile bands. (D) Minimum feasible travel time as a fraction of observed duration for the 25 zebrafish branches (post-epiboly); the *C. elegans* median (0.21) is marked. (E) Ensemble-floor calibration: per-branch observed action ratio versus the forward-simulated ensemble ratio (orange) and the random-partition ratio (gray) for the 25 zebrafish branches (log–log; dashed line: identity). Observed ratios sit at or below the ensemble floor and are indistinguishable from random partitions; the zebrafish excess is thus attributed to centroid geometry, not route suboptimality.

The calibrated headline measurement is therefore not the raw 3.79× ratio but the ≈2-fold path-specific excess of realized worm lineage paths over the model’s own ensembles; the zebrafish figure delimits where the measure reports ensemble geometry rather than route choice.

### 2.4 Cross-species replication: zebrafish embryogenesis at the ensemble floor

Are the action-based measurements specific to *C. elegans*, with its invariant, unusually deterministic lineage? We replicated the core LADD measurements on the zebrafish atlas (38,731 cells, 12 stages spanning 3.3–12 h post-fertilization (hpf); Fig 1E), generalizing the dynamics to per-step time increments for non-uniformly spaced stages (Methods 4.12). The learned field passed the same forecasting checks as in the worm (S3 and S10 Notes). We defined trajectories from the authors’ URD differentiation tree as 25 root-to-tip branch paths (per-stage centroids; Methods).

Over the full 3.3–12 hpf window, the action ratio was very large (median 310, IQR 248–336) but budget-sensitive: the minimum-action path rides the learned field through the epiboly burst, so modest differences in the achievable minimum swing the ratio (S10 Note). Regardless of budget, 78–95% of each path’s action was concentrated in the single OBLONG→DOME transition (3.8–4.3 hpf), where the centroid moves ∼15 principal-component units in 0.5 h under single-batch conditions (Methods; Fig 3C). The full-window ratio therefore measures the temporal burstiness of epiboly more than route choice, and we treat the post-epiboly window (4.7–12 hpf) as the primary comparison, matching the steady morphogenesis regime of the worm window.

In the post-epiboly window, zebrafish trajectories deviate from least-action paths by a median factor of 8.7 (IQR 7.8–10.0, n = 25 branches; Fig 3A), the same order of magnitude as the worm’s raw 3.9, and all controls reproduced the worm pattern (waypoint shuffling raised the action 10.3-fold, time reversal 4.3-fold; observed costlier than straight-line in 21/25 branches, median S_observed/S_line = 1.31; Fig 3B). Manifold-constrained and unconstrained minimum-action paths agreed (ρ = 0.995). The geometry-versus-timing decomposition recovers only a 1.7× factor from optimal re-timing, so most of the ≈8× excess is geometric rather than temporal. The bounded-control minimum-time analysis gave a median T_min/T_obs of 0.42 (IQR 0.36– 0.51; Fig 3D): every branch could connect its endpoints faster, but the margin is smaller than the worm’s 0.21 (Section 2.9), as expected once the epiboly burst is excluded.

The decisive difference is calibration: zebrafish observed ratios sit at the model’s forward-simulated floor (2.3; Fig 3E) and are indistinguishable from random partitions, so the zebrafish excess is fully accounted for by centroid geometry. But the raw magnitudes are directly comparable once temporal resolution is matched (Section 2.6).

One genuine cross-species difference emerged in temporal placement: the action-rate profile is *front*-loaded in zebrafish (pooled ρ = −0.85; 24/25 branches negative), driven by the epiboly burst, and remains so post-epiboly (ρ = −0.69, p = 2×10⁻²⁹). In the worm the profile is back-loaded (ρ = +0.54), peaking in the mid-embryonic fate-execution window (Fig 3C). The excess is thus of the same order across species over steady morphogenesis, while its temporal placement reflects each organism’s biology: zebrafish pays up front in epiboly, the worm late during fate execution.

### 2.5 Comparison with optimal-transport and flow-matching methods

We compared LADD’s learned dynamics with Waddington-OT (WOT) [2], CellRank [5] (moscot optimal-transport kernel [22]), and conditional flow matching (CFM) [4] on 12 held-out forecasting tasks per species (four start bins × three horizons; S7 Fig). All four methods beat the static baseline on all 24 tasks. The frameworks optimize different objectives, and each metric rewards its own: Optimal-transport couplings interpolate the observed marginals directly, which is what distribution-level accuracy (squared 2-Wasserstein distance to the held-out population) rewards, so the transport-based methods are strongest there (median relative W2: CellRank 0.25, WOT 0.47 in worm; 0.14 and 0.34 in zebrafish); LADD, which optimizes one-step dynamical consistency, is comparable to CFM (0.84 and 0.85 in worm; 0.63 and 0.64 in zebrafish) and more accurate than CFM in 9/12 tasks in both species. The same ordering holds at path level, where WOT’s barycentric projection interpolates the observed centroids by construction (S4 Note; S9C Fig). For the question this paper asks, the relevant quantity is the action functional, which the interpolative frameworks do not define.

A trajectory-level control confirms that the excess action is a property of the observed centroid paths, not of the learned field. We generated the trajectory implied by each method from the same starting points and scored them under the LADD action functional (Methods 4.15). A deterministic LADD rollout sits at the action floor under its own field (median ratio < 3×10⁻¹²), and an independently learned CFM rollout also remains near the floor (median 0.22× in worm, 2.8× in zebrafish; Wilcoxon against observed p = 4×10⁻³¹ and 6×10⁻⁸): the two fields agree on what a typical trajectory looks like. The CellRank barycenter, by contrast, carries an excess indistinguishable from the observed lineage centroids (median 3.9× vs 3.8× in worm, p = 0.17; 8.8× vs 7.9× in zebrafish, p = 0.47; S12 Note). This is the ensemble floor of Section 2.3 seen from an independent direction: a centroid of a spreading, bifurcating distribution departs from single-particle dynamics to the same degree, whether taken from data or from an optimal-transport pushforward.

### 2.6 Validity boundary I: the action ratio is resolution-dependent

The cross-species comparison rests on atlases with different temporal structures, so we quantified both ratios’ sensitivity to temporal resolution (Fig 4A,B). In the worm, the ratio is stable to interpolation (3.82 and 3.81 at 0.25- and 0.5-h grids vs 3.79 native) and declines gradually under bin merging (2.30 at 1 h, 1.58 at 1.5 h): the excess is not a fine-graining artifact, though very coarse grids erode it. In zebrafish, the full-window ratio is resolution-sensitive (310 native; 149–187 under interpolation), confirming that it measures burst timing, while the post-epiboly ratio is stable across 0.5–2-h grids (6.1–9.7) and equals 2.41 on a uniform 1-h grid over 7–12 hpf. The monotone decline under coarsening is the expected smoothing signature of centroid paths. Cross-species numbers must therefore be compared at matched resolution: on a 1-h grid the two organisms essentially coincide (worm 2.30 vs zebrafish 2.41), while each native-resolution figure (3.79 vs 8.7) reflects its own sampling density and dynamics range.

**Fig 4.**
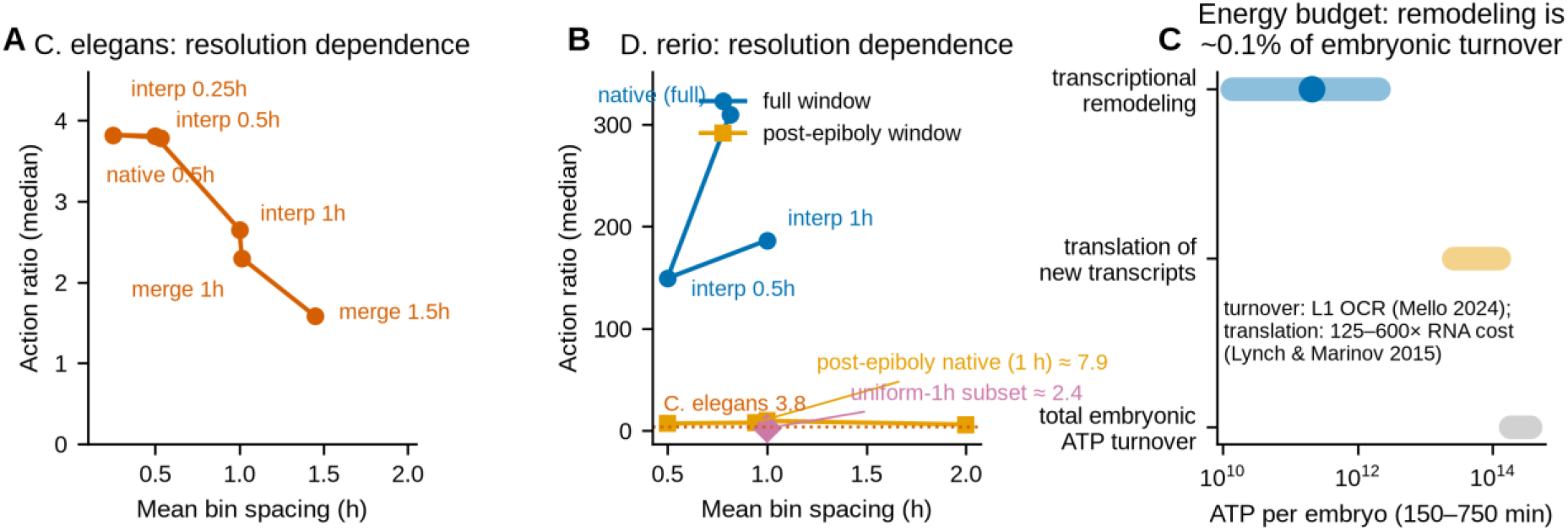
Validity boundaries of the action ratio. (A) *C. elegans*: median action ratio (179 paths) versus mean bin spacing under interpolation (0.25–2 h) and bin merging (1–1.5 h). The native 0.5-h grid gives 3.79. (B) Zebrafish: median branch ratio under interpolation and subsampling, for the full window (blue; dominated by the epiboly burst) and the post-epiboly / uniform-grid windows (orange; stable at 6–10). The uniform-1-h subset (purple diamond) gives 2.41; worm value marked for reference. (C) ATP equivalents per embryo over 150–750 min (log scale): transcriptional remodeling (blue; range: 5–20% detection efficiency, 2–48 ATP per nucleotide; point: central estimate), translation of the new transcripts (orange; 125–600× the RNA cost [23]), and total embryonic ATP turnover (gray; L1-anchored oxygen consumption [25], P/O ≈ 2.5). Transcriptional remodeling accounts for ≈0.1% of the budget.

### 2.7 Validity boundary II: the action is not a transcriptional energy cost

What, physically, does the excess action cost? Its rate should rise with active transcriptome remodeling, which carries a measurable energetic price. We translated the action-rate profile into adenosine triphosphate (ATP) equivalents using the atlas itself: newly synthesized transcripts per cell per 30-min bin, estimated from positive-part increments of per-gene raw counts between consecutive bin centroids (a conservative lower bound), converted with literature per-nucleotide costs [23] (Methods 4.19).

Two results emerged. **First, the action rate is coupled to transcriptional remodeling within lineages.** The per-path correlation between action rate and new-transcript rate was positive in 101 of 136 worm paths with ≥6 scored steps (median ρ = 0.36; sign test p = 1×10⁻⁸; Fig 5A,C) and in all 25 zebrafish branches (median ρ = 0.71, p = 6×10⁻⁸; Fig 5C). The action rate thus partly tracks the pace of gene-expression change, as a cost reading predicts. **Second, the transcriptional cost is negligible against the embryonic energy budget.** Summing per-cell synthesis over the embryo’s cell-count curve gives ≈9×10⁵ detected new transcripts embryo-wide over 150–750 min; at 5–20% detection efficiency [24] and 2–48 ATP equivalents per nucleotide [23], transcriptome remodeling costs ≈10¹⁰–10¹² ATP equivalents per embryo, against a total embryonic ATP turnover of ≈2–4×10¹⁴ (L1-anchored oxygen consumption [25], phosphate/oxygen [P/O] ratio ≈ 2.5). The remodeling cost is therefore ≈0.1% of the budget (range 0.004–1%; Fig 4C); downstream translation of the new transcripts can reach 10–50% of turnover (S11 Note). The remodeling rate also does not track whole-embryo metabolism where measurable: in zebrafish, the mean transcription rate falls over 5–12 hpf while oxygen consumption rises (profile-shape ρ = −0.52, not significant; Fig 5B) [26]. The embryo’s energy budget buys morphogenesis (biomass, membranes, division [27]) while transcriptomic speed costs almost none of it; but because translation-coupled costs reach 10–50% of turnover, energetics cannot be excluded as the selective pressure a least-action principle would answer to. Whether the action is better read as a control or information cost remains an open question.

**Fig 5.**
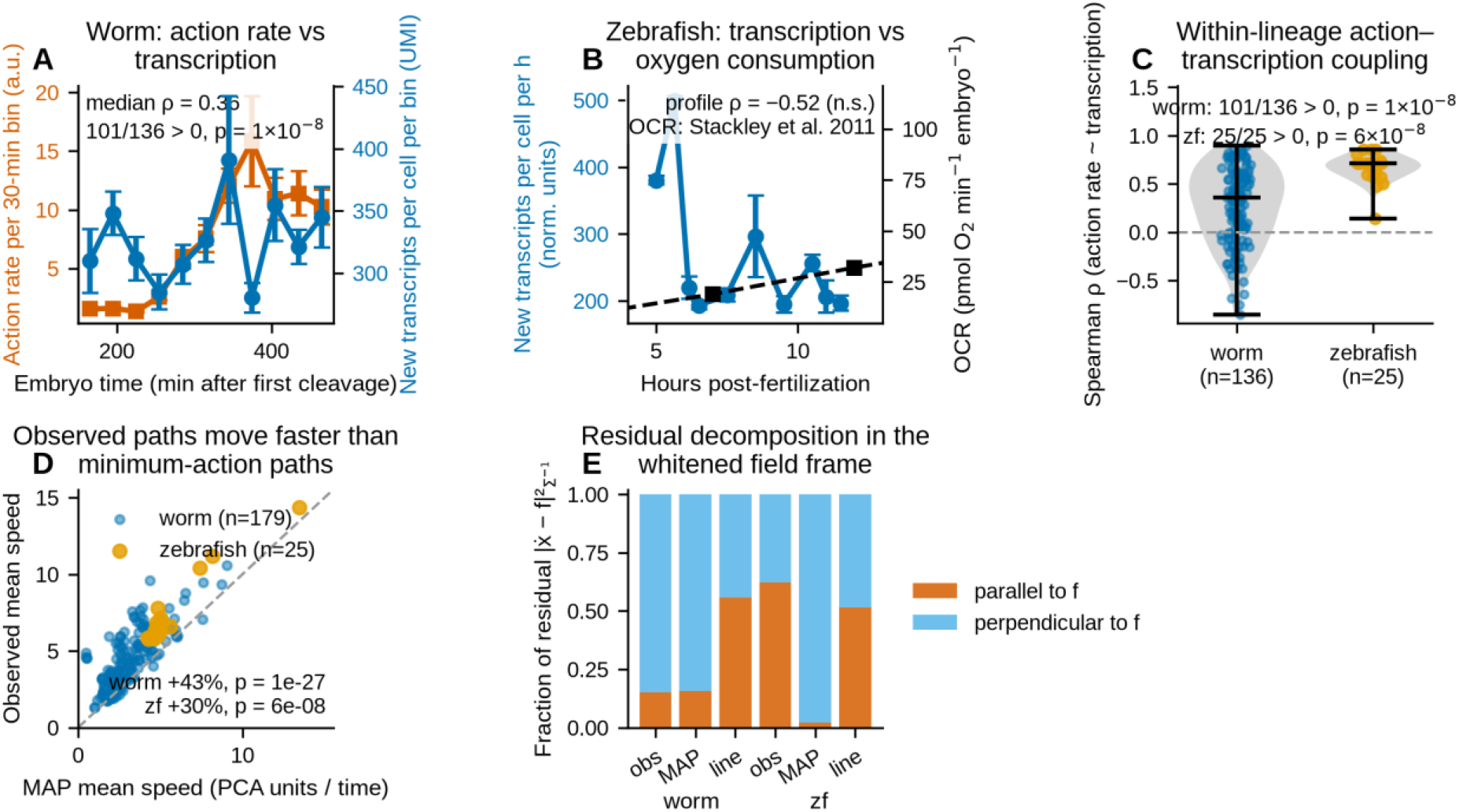
Mechanistic readings of the action: transcriptional coupling and the speed test. (A) Worm mean action rate (vermillion, left axis) and mean transcriptional-remodeling rate (blue, right axis; new transcripts per cell per 30-min bin, positive-part estimator) across lineage paths (mean ± standard error; bins with ≥20 contributing paths). (B) Zebrafish mean transcriptional-remodeling rate (blue, left axis; normalized-count units per hour) across URD branches, with measured whole-embryo oxygen consumption (black dashed; digitized from Stackley et al. [26]). The profiles are anti-phasic (Spearman ρ = −0.52, not significant). (C) Within-lineage Spearman coefficients between action rate and transcription rate in worm (n = 136; median ρ = 0.36; 101/136 positive, sign test p = 1×10⁻⁸) and zebrafish (n = 25; median ρ = 0.71; 25/25 positive, p = 6×10⁻⁸). (D) Mean traversal speed of observed versus minimum-action paths (worm +43%, Wilcoxon p = 1×10⁻²⁷; zebrafish +30%, p = 6×10⁻⁸; dashed line: equality). (E) Residual ẋ − f decomposed into field-parallel (vermillion) and field-perpendicular (sky) fractions in the Σ⁻¹-whitened frame for observed, minimum-action, and straight-line paths in both species.

### 2.8 The calibrated excess action tracks path geometry, not fate-map complexity

With the instrument calibrated and its validity boundaries stated, we return to the worm’s path-specific excess and ask what it reflects. A natural hypothesis is that the excess action is the price of fate diversification: lineages with more fate decisions or descendant cell types should pay more. We mapped all 179 branch anchors onto the reference embryonic lineage tree [28] and computed descendant and branch-point counts per subtree, atlas-based diversity (rarefied cell-type and subtype counts), and window-matched diversification (Methods).

Raw correlations appeared to support it (descendant count: Spearman ρ = 0.40, p = 3.6×10⁻⁸; rarefied cell types: ρ = 0.49, p = 3.7×10⁻¹²; S5A,C Fig; full statistics in S8 Note). But fate complexity is deeply confounded: large subtrees are shallower, longer-lived, and more densely sampled, and these are the same geometric factors that mechanically inflate the ratio. Controlling for depth, path duration, and sampling by partial rank correlation abolished every fate-complexity association (partial ρ between −0.13 and +0.14; all q > 0.15; S5B,C Fig), and 40 sister-branch pairs differing in descendant count showed no tendency for the more complex sister to carry higher excess (52.5% concordant, binomial p = 0.87; S5D Fig). Under the same controls, path geometry does predict the excess: curvature (partial ρ = 0.46, p = 1.8×10⁻¹⁰; S5F Fig) and duration (partial ρ = 0.41, p = 1.1×10⁻⁸). The depth and temporal gradients of Section 2.2 are themselves abolished under these controls: the ratio measures geometric and temporal structure, not fate-map cost.

Finally, our trajectories are branch *centroid* paths, which smooth single-cell heterogeneity. Lineage-consistent single-cell chains (168 anchors; Methods) exceeded the corresponding centroid-path action in 88% of cases (median 1.9×, IQR 1.4–2.6; Wilcoxon p = 7.6×10⁻¹⁶; S5E Fig). Because the calibration floor (2.3) is defined for the same centroid ensembles, all ratios here remain internally consistent centroid-path quantities.

### 2.9 The time margin of development

A complementary question is the brachistochrone one: how much faster *could* development be? For 30 lineage anchors stratified by depth, we computed the minimum feasible travel time T_min between each trajectory’s observed endpoints under the controlled dynamics dx = (f + u)dt, bounding the steering magnitude by the peak the observed trajectory used (|u| ≤ u_max; Methods). The minimum feasible time was a median 21% of the observed duration (T_min/T_obs = 0.21, IQR 0.16–0.27; S8F Fig): lineages take roughly five times longer to connect their endpoints than their steering envelope requires. The margin agrees with the action ratio (Spearman ρ = −0.74, p = 3×10⁻⁶) and is largest for bursty lineages whose peak steering far exceeds their average. The action–duration frontier quantifies the speed–cost trade-off: halving traversal duration multiplies the minimum action ≈1.6-fold (S8G Fig). Because the feasibility optimization is not manifold-constrained, these margins are upper bounds.

### 2.10 Observed paths outrun the learned field rather than fight it

The decomposition of Section 2.2 assigns 87% (worm) of the log-excess action to the geometric factor, yet set-level route comparisons find minimum-action paths no closer to the observed route than a straight line, so the geometric share cannot be read as spatial detour. Because re-timing holds the curve fixed, the geometric factor absorbs any deviation that is not pure schedule along it, including traversal-speed differences. We therefore tested directly what observed paths do differently: along each observed, minimum-action, and straight-line path we evaluated the field magnitude |f|, the signed field projection on the direction of motion, and a metric-consistent decomposition of the residual ẋ − f (Methods 4.20).

One possibility is that observed paths fight the flow or exploit strong-field regions. The data reject this in both species. Observed paths traverse slightly *weaker*-field regions than minimum-action paths (median |f| 1.80 vs 1.92 in worm, p = 8×10⁻⁹; 3.77 vs 4.02 in zebrafish, p = 5×10⁻⁵), and they do not move against the flow: the signed field projection is positive on average, indistinguishable from minimum-action paths in worm (p = 0.67) and significantly *greater* in zebrafish (p = 4×10⁻⁴). Straight-line routes, by contrast, move against the field in 93% (worm) and 65% (zebrafish) of steps, so the learned flow itself opposes endpoint shortcuts. What distinguishes the observed paths is speed: they traverse state space 43% (worm, p = 1×10⁻²⁷) and 30% (zebrafish, p = 6×10⁻⁸) faster than minimum-action paths between the same endpoints (Fig 5D), consistent with the timing factor R_time > 1 (2.2) and the minimum-time margin (2.9). The residual decomposition adds a nuance: in the worm, observed and minimum-action paths carry the same residual structure (∼85% field-perpendicular meandering; p = 0.17), while in zebrafish the observed residual is 62% field-parallel (speed modulation along the flow), against 2% for minimum-action paths (p = 6×10⁻⁸; Fig 5E). The excess action is therefore not paid to fight the dynamics or to exploit fast regions. It is the price of moving faster than, and off, the least-action route.

### 2.11 The excess action couples to morphogenetic events and cell-cycle slowing

We bundled three pre-specified tests with permutation nulls and Benjamini–Hochberg correction (Fig 6A,B). (H1) Event alignment: action is significantly concentrated in the pharynx-morphogenesis window (330–430 min: observed 27.9% vs duration-matched circular-permutation null 15.3%, p = 10⁻³, q = 0.005) and in the union 330–480 min fate-execution window (30.7% vs 19.5%, q = 0.005), but not in the muscle, enclosure, or neural windows (all q > 0.3). The alignment is specific, not a generic mid-developmental artifact. (H2) Division-rate coupling: within each path, the per-step action rate anti-correlates with the local division rate (median Spearman ρ = −0.29; 34/179 paths positive; sign test p = 1.6×10⁻¹⁷). (H3) Cell-cycle slowing: the action-rate profile correlates positively with the literature cell-cycle-length curve over 150–720 min (median ρ = +0.66; 159/179 positive; p = 4.6×10⁻²⁸). High-action segments therefore coincide with slow-cycling, morphogenesis-executing periods.

**Fig 6.**
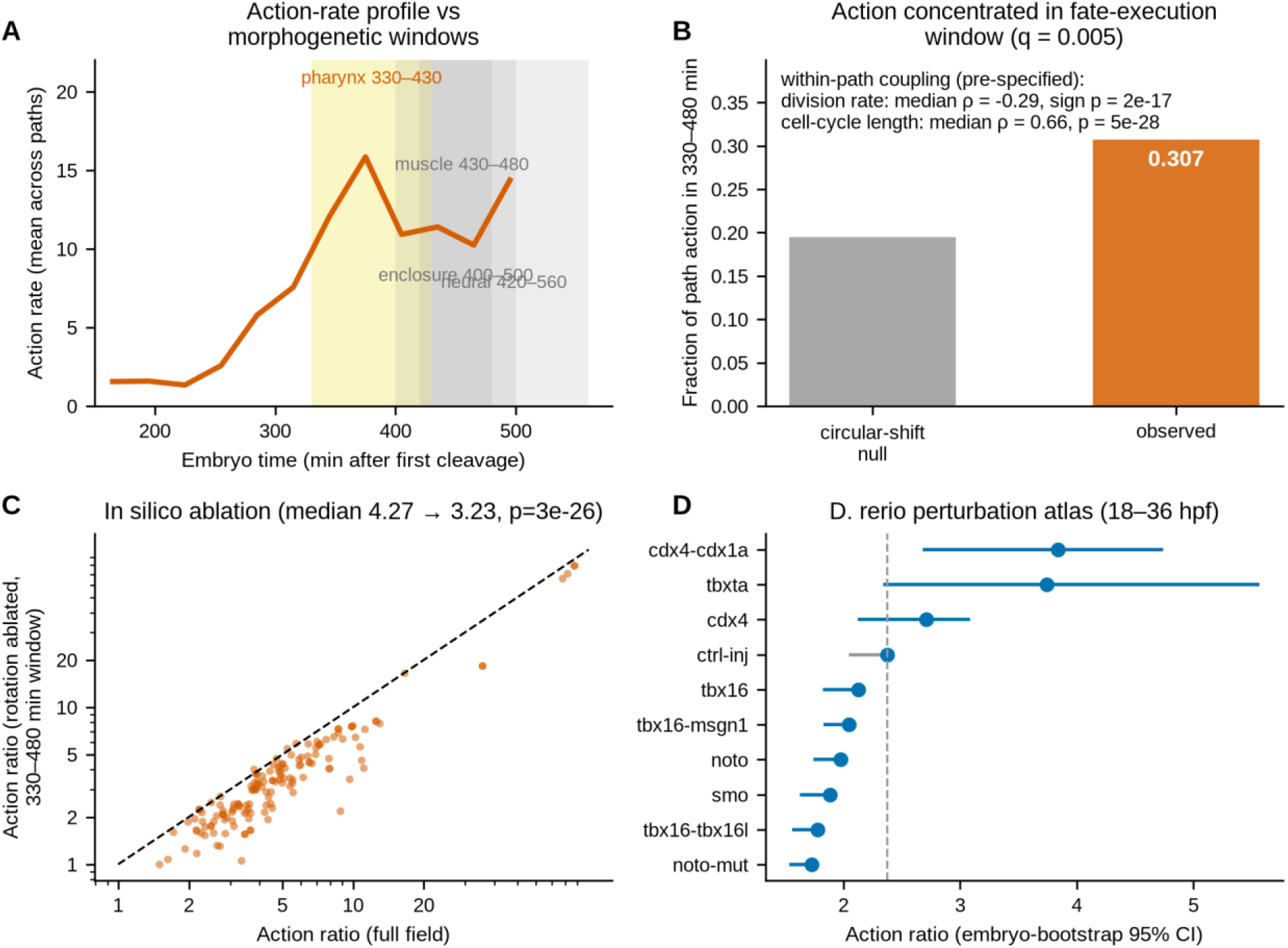
The excess action couples to morphogenesis and responds to perturbation. (A) Mean per-step action rate across the 179 worm paths over developmental time. Shaded bands: literature morphogenetic windows (yellow: pharynx 330–430 min, significant; gray: muscle, enclosure, neural, not significant). (B) Fraction of path action in the 330–480 min window, observed versus circular-permutation null (30.7% vs 19.5%, p = 10⁻³, q = 0.005; pharynx sub-window 27.9% vs 15.3%, q = 0.005). Within-path coupling: action rate anti-correlates with division rate (median ρ = −0.29, sign test p = 1.6×10⁻¹⁷) and correlates with the cell-cycle-length curve (median ρ = +0.66, p = 4.6×10⁻²⁸). (C) In silico ablation of the rotational field component within 330–480 min: per-path action ratios before and after (log–log; dashed line: identity; median 4.27 → 3.23, Wilcoxon p = 3×10⁻²⁶, n = 152 paths). (D) Zebrafish perturbation atlas: action ratios under the control field for ctrl-inj and nine perturbation conditions (points: estimate; bars: embryo-bootstrap 95% CI; dashed line: control).

### 2.12 Perturbations move the action ratio in the expected directions

If the excess action reflects active steering, disrupting patterning should change it. We tested this in silico and in real perturbation data (Fig 6C,D). (i) In silico ablation: removing the rotational (non-gradient) field component specifically within the 330–480 min window, a component elevated at fate-decision nodes (S6 Note), reduced the median ratio from 4.27 to 3.23 across the 152 paths traversing the window (Wilcoxon p = 3×10⁻²⁶). Observed-path action is nearly unchanged (0.997×), but the minimum achievable action rises 1.36×: the window’s non-gradient steering makes observed paths deviate from a purely gradient-like flow. (ii) Temporal rescaling of the window changes the ratio monotonically (median 5.9–14.4 across rescale factors 0.5–2.0; S8H Fig), so the excess reflects realized duration, not a fixed geometric penalty. (iii) Real perturbations: we applied LADD to the zebrafish perturbation atlas of Saunders et al. [29] (240,000 cells; 18/24/36 hpf), learning a control field from ctrl-inj embryos and scoring condition-pooled centroid paths (embryo-level bootstrap intervals). The control ratio is 2.37 [2.05, 2.40], and seven of nine perturbations significantly shift it (p < 0.05): cdx4;cdx1a double mutants increase it (3.84 [2.68, 4.74], p = 0.006), while tbx16-family, noto, and smo perturbations decrease it (e.g., tbx16;tbx16l 1.78, p < 10⁻⁷; noto-mut 1.73, p < 10⁻⁹; full statistics in S9 Note). Restricting paths to mesoderm cells preserves the pattern in the affected germ layer (cdx4: 2.48 vs mesoderm control 1.68, p = 4×10⁻⁵). Mesoderm- and axis-patterning perturbations therefore move the ratio in both directions, supporting it as a functional readout of program integrity, not an atlas constant.

## 3. Discussion

Single-cell atlases now infer developmental dynamics from destructive snapshots, and every use of those dynamics (prediction, landscape analysis, optimality comparison) inherits the construction. We asked how far realized lineage paths depart from the routes such inferred dynamics consider natural, and how much of the apparent departure is the measurement itself. Variational principles are increasingly imported into developmental biology, sometimes as the explicit claim that development follows minimum-action paths [10]; atlases make such claims computable. Our answer has two parts. The first is the instrument: LADD learns a stochastic developmental vector field from time-series scRNA-seq and computes minimum-action and minimum-time paths comparable, with controls, to realized trajectories. The second, the central contribution, is the calibration: raw optimality ratios are dominated by a generic measurement cost, and three calibration checks (configuration stability, circularity controls, null-ensemble floors) separate this cost from path-specific deviation. The headline number is therefore not the raw worm ratio of 3.79× but the calibrated one: realized lineage paths carry a ≈2-fold path-specific excess over the model’s own ensembles (median fold 1.96, p = 4×10⁻³¹), while the zebrafish excess sits entirely at the ensemble floor. Without calibration, the same pipeline would report “development is ≈4–9× suboptimal” in both species: mostly artifact in one, entirely artifact in the other.

The calibrated excess is specific in composition. It is geometric before it is temporal (87% of the log-excess in worm, 74% in zebrafish), yet not produced by detours against the dynamics: observed paths move 30–43% *faster* than minimum-action paths, through weaker-field regions, aligned with the learned flow. Nor is it the price of fate diversification: the pre-specified confounder analysis returned a clean null for every fate-complexity metric once path duration, lineage depth, and sampling density were controlled. The excess is also biologically responsive: it concentrates in the mid-embryonic fate-execution window, tracks cell-cycle slowing, drops when the field’s non-gradient component is ablated in silico, and shifts in both directions under zebrafish patterning perturbations. The minimum-time analysis adds a complementary bound: even granted its peak steering effort as the speed limit, development uses only about a fifth of its attainable pace, a statement about the speed–cost trade-off, not slowness.

The calibrated claim is a measurement, not a mechanism: it establishes a path-specific excess and localizes its immediate correlate to speed (detours, flow-opposition, and fate complexity are excluded), while the function of the excess (selected or tolerated) and its single-cell resolution remain open questions for non-destructive profiling.

The measurement’s validity boundaries are as much a deliverable as the numbers. The action ratio is resolution-dependent: it survives interpolation but erodes under bin merging, so cross-species comparisons are meaningful only at matched temporal resolution, where the worm and zebrafish values coincide (2.30 vs 2.41 at 1 h). The action is also not a transcriptional energy cost: the remodeling coupled to the action rate is ≈0.1% of the embryonic ATP budget, and where respiration can be measured it runs anti-phasic to remodeling; but translation-coupled costs reach 10–50% of turnover, so energetics overall cannot be excluded. Whether the action is better read as a control or information cost is speculation we state without measurement support.

### Limitations

(i) The action is defined relative to the *learned* autonomous dynamics in a 50-dimensional transcriptomic latent space; a proxy for organismal fitness or energetic cost, not a measure. A time-conditioned field f(x, t) leaves all conclusions unchanged (2.3). (ii) Lineage trajectories are branch *centroid* paths (destructive scRNA-seq measures each cell once); single-cell chains carry ≈1.9× higher actions, but the calibration floor is defined for the same centroid ensembles, keeping ratios consistent. (iii) The latest time bins are moderately batch-dominated, and lineage coverage falls after ∼550 min, restricting analysis to the fate-specification window. (iv) The manifold-constrained optimization may not find the global optimum, and the worm holdout is limited to 13 fully held-out deep paths; the action ratio is an upper bound on suboptimality. The zebrafish speed analysis (2.10) uses paths recomputed at a higher optimization budget than in 2.4 (residual sensitivity: median 7%, max 27%, EVL/periderm). (v) The mechanism-association analysis is correlational and restricted to 179 overlapping branch anchors; fate complexity is intrinsically confounded with path geometry, so causal claims require perturbation experiments. (vi) The cross-species comparison is order-of-magnitude except at matched resolution; the zebrafish atlas has stage-level time resolution, URD branches rather than invariant-lineage paths, and single-batch early stages, hence the post-epiboly window. The perturbation analysis (2.12) uses condition-pooled centroid paths over 18–36 hpf from an atlas not overlapping the Farrell et al. window; ratios are compared to their own control, not to developmental-atlas values. (vii) The action ratio grows with latent-space dimensionality (≈3.2 in 20-D, 3.8 in 50-D, 5.0–5.5 in 100-D PCA); the 50-D primary choice balances action-independent criteria (2.3), and qualitative conclusions are unchanged. (viii) The energetic translation (2.7) is an order-of-magnitude calibration, not a thermodynamic identity (24-fold per-nucleotide cost range; 5–20% detection efficiency; lower-bound synthesis estimator; L1-anchored worm turnover).

### Outlook

For the growing literature on variational readings of development, an action ratio without a calibration floor is uninterpretable: the same number can mean route suboptimality, ensemble spread, temporal burstiness, or resolution mismatch. The calibrated claim is narrower than any raw optimality ratio, and, we argue, more interesting. The framework is organism-agnostic; frog and mouse atlases are natural next steps, provided comparisons use matched resolution against calibrated floors. Two extensions would connect kinematics to mechanism: time- and lineage-conditioned dynamics, and direct, time-resolved measurements of embryonic metabolism. More broadly, LADD shows one way to import a physics variational principle into biology: not as an assumed law, but as a falsifiable measuring instrument with a stated noise floor.

## 4. Methods

### 4.1 Data

We used the lineage-resolved *C. elegans* embryogenesis single-cell atlas of Packer et al. [20] (GEO GSE126954; processed AnnData from Zenodo record 15236812, file ‘celegans_global_adata.h5ad’, MD5-verified). Of 96,084 cells, we retained 89,701 that passed the authors’ quality-control flag (‘to.filter == FALSÈ). We then restricted the data to 150–750 min after first cleavage (the authors’ ‘embryo.timè coordinate; 87,702 cells), where sampling density was adequate. Annotations used: ‘embryo.timè (continuous, min), ‘lineagè (invariant-lineage position), ‘cell.typè, ‘batch’.

### 4.2 Preprocessing

We normalized counts to 10,000 per cell and applied log1p. We selected 3,000 highly variable genes (Seurat method). We scaled the data (clipped at ±10) and reduced it to 50 principal components (29.9% cumulative variance). We assessed batch–time confounding by the maximum single-batch fraction per 30-min bin. We excluded lineage strings containing an ambiguous multi-candidate separator (“/”) or marked “unannotated” from lineage analyses. We retained the “x” token (unresolved division) as a node label. This yielded 31,753 lineage-usable cells across 368 distinct lineage annotations.

### 4.3 LADD: dynamics learning

We model latent dynamics as dx = f_θ(x)dt + Σ^(1/2)dW. The drift f_θ is a multilayer perceptron (MLP; 50→256→256→256→50, SiLU). We trained it to match empirical one-step displacements between adjacent 30-min bins. For each adjacent bin pair, we estimated each source cell’s displacement as the mean displacement to its k = 10 nearest neighbors in the next bin (local coupling). This gives a robust estimate of the conditional mean displacement. We trained the drift by minimizing the mean squared one-step Euler error (AdamW, lr 10⁻³, cosine schedule, 200 epochs, batch 4096). We calibrated the diffusion covariance Σ as the time-average of the positive-semidefinite-projected difference between the observed next-bin covariance and the deterministic-rollout covariance. We also evaluated global entropic optimal-transport coupling and rejected it, because its blurred couplings produced divergent rollouts (Supplementary Note). For landscape visualization, we fitted a scalar potential U(x) (MLP 50→256→256→1) to 30,000 subsampled cells by minimizing ||−∇U(x) − f(x)||² (AdamW, lr 10⁻³, 200 epochs). We computed the per-cell rotational fraction as ||f + ∇U||²/||f||² on 20,000 subsampled cells.

### 4.4 Forecasting validation

Leave-one-time-bin-out (LOBO) interpolation: for each held-out bin b in the evaluable interior range (bins 4–16, i.e., bin centers 285–645 min; bins outside this range lack sufficient cells or a usable predecessor transition), we retrained the drift on all other transitions and used it to predict the held-out bin from its predecessor. Forward extrapolation: we trained the drift on all transitions up to a cutoff (bins 6, 9, 12, 15) and rolled it out 1–3 bins (30–90 min) into unseen times. We compared predicted and observed distributions by squared 2-Wasserstein distance (exact earth-mover’s-distance computation on 3,000-cell subsamples) and radial-basis-function (RBF) kernel maximum mean discrepancy (MMD). Baselines: persistence (no change) and ridge-regression linear drift trained on the same pairs.

### 4.5 Lineage-based validation

Branch trajectories: for each lineage anchor prefix, we took the per-bin centroid of cells whose lineage string starts with that prefix (≥10 cells/bin, ≥6 bins). This gave 179 centroid paths. Fate prediction: we rolled cells forward and assigned them the nearest terminal-type centroid (types with ≥20 annotated cells at 550–700 min). We scored balanced accuracy against the authors’ ‘cell.typè annotation, and we computed per-type recall for the 390→510 min rollout. Lineage coherence: we compared sister lineages (differing only in the terminal character) to random pairs by the distance between their rolled-out futures (Mann–Whitney U). For the relatedness-decay analysis, we rolled lineage centroids at bins 4, 6, and 8 (≥3 cells) forward by 3 bins. We binned all within-founder pairs by shared path depth in the reference lineage tree (maximum longest-common-prefix length over wildcard-consistent node matches). We then computed the Spearman correlation between shared depth and predicted future distance over 9,454 pairs.

### 4.6 Action analysis

The Freidlin–Wentzell action of a discrete path x(t) is S = ½∫|ẋ − f(x)|²_Σ⁻¹ dt. We computed minimum-action paths between fixed endpoints over the same duration by direct collocation (Adam on free interior waypoints, 300–400 iterations). We initialized from both a straight line and the observed path, and we retained the lower action. Controls: time-reversed path; waypoint-shuffled path (5 permutations); straight-line path. To prevent off-manifold shortcuts, we also computed minimum-action paths under a k-NN-distance manifold penalty (k = 5) anchored to the observed cell density. We quantified route similarity by the symmetrized Chamfer distance between the minimum-action path and the observed path, relative to a straight line.

Geometry-versus-timing decomposition: for a fixed geometric curve, the action is a sum of per-segment terms g_i(Δt_i) = ½(a_i/Δt_i − 2b_i + c_iΔt_i) with a_i = |Δx_i|²_Σ⁻¹, b_i = ⟨Δx_i, f(x_i)⟩_Σ⁻¹, c_i = |f(x_i)|²_Σ⁻¹. Each term is convex in the segment time Δt_i. The exact optimal re-timing under the constraint ΣΔt_i = T follows from the KKT condition Δt_i(λ) = √[a_i / max(c_i – 2λ, ε)], with λ found by bisection such that the segment times sum to T. R_time = S_obs/S_retimed and R_geo = S_retimed/S_min then multiply to R_obs exactly. As a secondary check, we reparameterized the observed curves at constant speed (uniform arc length in the Σ⁻¹ metric, same waypoint count and total duration) and rescored them.

### 4.7 Reference lineage tree and fate-complexity metrics

We used a machine-readable encoding of the Sulston reference embryonic lineage [28] (2,189 nodes; 1,444 embryonic nodes after excluding post-embryonic descendants). For each branch anchor, we counted the terminal descendants and branch points (nodes with ≥2 embryonic children) in the anchor’s subtree. We mapped atlas lineage strings containing unresolved divisions (“x”) to reference nodes by wildcard matching over the Sulston division symbols {a, p, l, d, r, v}, with progressive prefix shortening when an anchor exceeded tree depth. We mapped single-letter pseudo-anchors via the longest common prefix of the matching atlas lineages. All 179 anchors mapped (173 at full length), and founder subtree counts matched canonical values (e.g., E = 20 intestinal cells). We quantified atlas-based diversity by rarefaction: we subsampled each anchor’s descendant cells to 200 cells (100 replicates) and recorded the mean numbers of distinct ‘cell.typè and ‘cell.subtypè annotations. Window-matched diversification metrics counted the cell types present at a path’s start and end bins (types gained) and the growth in annotated terminal lineages over the path window.

### 4.8 Mechanism-association statistics

We assessed associations between the action ratio and each complexity metric by Spearman correlation. For confounder-controlled associations, we used partial Spearman correlation on rank residuals. We residualized ranked variables on ranked lineage depth, path duration in bins, and log₁₀ descendant cell count by ordinary least squares, then computed the Pearson correlation of residuals (t-test with n−k−2 degrees of freedom). We adjusted p-values by Benjamini–Hochberg across metrics. Sister-subtree pairs were anchor pairs differing only in the terminal division symbol, with both members present and with unequal descendant counts. We tested concordance (the higher-complexity sister carries the higher action ratio) by an exact binomial test and a Wilcoxon signed-rank test. Mean log₁₀ unique molecular identifier (UMI) count per anchor served as a negative-control metric.

### 4.9 Single-cell lineage chains

For each anchor with ≥5 candidate start cells and a path spanning ≥4 bins, we reconstructed five single-cell chains by greedy nearest-descendant stitching. We started from a random cell of the anchor group in the first bin. Each subsequent bin contributed the nearest cell (PCA space) whose lineage string extends the current cell’s lineage. When no direct descendant was sampled, we fell back to the anchor group. We computed chain actions exactly as for centroid paths, and we compared each anchor’s median chain action to its centroid-path action (Wilcoxon signed-rank test).

### 4.10 Action-rate profiles and rotational localization

We computed per-step action rates along each observed centroid path as ½|Δx − f(x)|²_Σ⁻¹ per 30-min step. We normalized within-path profiles to each path’s mean rate and aligned them by normalized path position (0 = start, 1 = end). We quantified the position–rate association by Spearman correlation over all steps of all 179 paths. We checked robustness by recomputing rates with per-bin calibrated diagonal diffusion. We computed the per-cell rotational fraction ||f(x) + ∇U(x)||²/||f(x)||² by automatic differentiation through the fitted potential. We designated reference-tree nodes with ≥2 embryonic children as fate-decision (branch-point) nodes. We mapped lineage-annotated cells to reference nodes as in 4.7, and we compared rotational fractions of cells at branch-point versus transit nodes within 60-min windows, with stratification by quintiles of |f(x)|.

### 4.11 Minimum-time analysis

For 30 anchors stratified by depth (10 founder, 10 mid, 10 deep; largest cell counts), we computed the minimum feasible travel time between each observed trajectory’s endpoints. We used the controlled dynamics dx = (f(x) + u)dt, with per-step steering magnitude bounded by the peak the observed trajectory itself used: u_max = max_i |Δx_i − f(x_i)| (Euclidean norm in PCA space, per 30-min model-time unit). We tested the feasibility of a candidate duration T by direct collocation (40 waypoints, 300 Adam iterations), minimizing the mean squared violation of the control bound, from both straight-line and observed-path initializations. We deemed T feasible if the optimized peak |u| ≤ 1.05·u_max. We located the minimum feasible T by bisection (6 iterations) over [0.05, 1.0]×T_obs. The feasibility optimization is unconstrained by the data manifold, so off-manifold shortcuts can only make T_min smaller. The reported time margins are therefore upper bounds on attainable speed-up. We computed the action–duration frontier as the manifold-constrained minimum action (4.6) at durations {0.5, 0.65, 0.8, 1.0, 1.3}×T_obs.

### 4.12 Zebrafish validation

We replicated the core measurements on the zebrafish embryogenesis atlas of Farrell et al. [21] (GEO GSE106474, Drop-seq subseries of GSE106587; the authors’ URD trajectory object from the Broad Single Cell Portal, SCP162). We restricted UMI counts (23,974 genes) to the authors’ filtered set of 38,731 cells with the authors’ stage annotations (12 stages, 3.3–12 hpf; hours-post-fertilization values taken from the authors’ metadata). Preprocessing mirrored the worm pipeline (10,000-count normalization, log1p, 3,000 highly variable genes, scaling, 50 principal components; 26.4% cumulative variance). The 12 stages are non-uniformly spaced in developmental time (intervals 0.4–1.0 h). We therefore trained the drift on k-NN-coupled pairs (k = 10) with per-pair time increments, normalized the diffusion-covariance calibration by the corresponding interval per stage pair, and used per-step increments in all action integrals; the model time unit is hours. Forecasting validation used leave-one-stage-out interpolation (10 interior stages) and forward extrapolation (training cutoffs at stages 5, 7, and 9; horizons of 1–2 stages) against persistence and a ridge-regression velocity baseline, with the same Wasserstein and MMD metrics as in 4.4. We defined developmental trajectories from the authors’ URD tree. For each of the 25 named terminal cell types, we extracted the root-to-tip chain of tree segments. The per-stage centroid of the chain’s cells (≥10 cells per stage, ≥6 stages; all 25 chains passed, spanning 9–12 stages) gave the observed path. Action analysis, controls (time reversal, waypoint shuffle, straight line), minimum-action paths (unconstrained and manifold-constrained, two initializations), and the bounded-control minimum-time analysis were computed exactly as for the worm (4.6, 4.11), with durations in hours.

### 4.13 Error propagation and model-configuration stability

Seventeen configurations were evaluated: the reference architecture (50-D latent, 256×3 hidden) at five seeds; 128×2 and 128×3 architectures at three seeds each; and 20-D and 100-D PCA latent spaces (reference architecture) at three seeds each. For each configuration, we retrained the field and recomputed all 179 path ratios. We obtained confidence intervals by nested bootstrap (2,000 replicates resampling configurations, then paths within configuration).

Action-independent dimensionality criterion: for the 20-, 50-, and 100-D latent spaces (three seeds each, reference architecture), we computed two criteria that do not involve the action. Forecasting skill: relative W2 (predicted over static) on the twelve worm forecasting tasks of 4.15. Lineage coherence: sister-versus-random separation of rolled-out futures (three-step rollout from the 330-min bin; 400 pairs each), as in 4.5. Variance explained was computed on the same z-scored highly-variable-gene matrix (22.2%, 30.0%, and 36.6% at 20-, 50-, and 100-D).

### 4.14 Circularity controls

Lineage holdout (worm): we split terminal lineages into five folds. For each fold, we retrained the field without any cell belonging to those lineages, and we scored branch paths whose entire lineage was held out (13 deep paths). Leave-one-branch-out (zebrafish): for each of the 25 URD branches, we retrained the field without that branch’s cells and recomputed the branch ratio. Model-agnostic drift: we fitted a global linear drift (ridge regression on k-NN displacement pairs) and a Gaussian-process drift (RBF kernel, 2,000 inducing points, maximum-a-posteriori hyperparameters) to the same training pairs. We recomputed actions and minimum-action paths under each surrogate for all 179 worm paths. Time-conditioned field: we extended the drift MLP with a sinusoidal time embedding and retrained it. We recomputed forecasting, path actions, and minimum-action paths with time-dependent drift.

### 4.15 Benchmark against WOT, CellRank, and CFM

We scored twelve forecasting tasks per species (start bins {3, 6, 9, 12} worm / stage indices {2, 4, 6, 8} zebrafish; horizons 1–3 bins) by squared 2-Wasserstein distance between predicted and held-out cell populations in PCA space. WOT (v1.0, ‘OTModel’ with local_pca=0) computed transport maps between adjacent bins; multi-step predictions composed the maps. CellRank used a moscot TemporalProblem (sequential policy, joint PCA space) with the RealTimeKernel transition matrix. CFM used torchcfm conditional flow matching trained on the same k-NN displacement pairs as LADD, integrated with a fixed-step ordinary-differential-equation (ODE) solver. The static baseline carried the source population forward unchanged.

For the trajectory-level benchmark, we regenerated each branch path of the main action analysis (179 worm paths at native 30-min resolution; 25 zebrafish paths in the post-epiboly window) from the observed start point. We used three procedures. (i) A deterministic Euler rollout of the LADD field with the per-step time increments of the observed path. (ii) Integration of the trained CFM ODE (40 Euler substeps per observed interval), sampled at the observed time points. (iii) Barycentric pullback through the CellRank transition matrix: a uniform distribution over the 30 nearest cells of the start bin is propagated through the row-stochastic transition matrix, and the trajectory position at each bin is the barycenter of the pushed-forward distribution. We scored all trajectories under the LADD action functional with the per-step time increments and the full residual covariance of the main analysis, and we normalized them by the per-path constrained minimum action. We compared ratio distributions against the observed paths by two-sided Wilcoxon signed-rank test.

Path-level one-step control: for every consecutive-bin step of the 179 observed worm centroid paths (1,310 steps), we predicted the next observed centroid three ways: persistence (the current centroid), a LADD ensemble rollout (one Euler step of the field from each of the lineage’s current-bin cells, then averaged), and a WOT barycentric projection (the lineage’s current-bin cells pushed through the row-normalized transport map above, then averaged). Errors are Euclidean distances to the observed next centroid in PCA space; per-path mean errors were compared by two-sided Wilcoxon signed-rank test.

Null-model calibration of the action functional: three controls bound the ratio’s ensemble floor. (i) Synthetic scenarios: we simulated 5,000 particles × 12 bins in 50 dimensions under four exactly solvable dynamics: static isotropic diffusion (no drift), coherent ballistic translation, a state-encoded 60/40 bifurcation (a persistent competence coordinate drives lateral lobe velocities from bin 4), and an asynchronous logistic wave (autonomous velocity field v(x) = x(8 − x)/6.4 with particles spread along the wave). We analyzed them by the identical pipeline (k-NN coupling, VectorField training, full-covariance calibration, centroid paths, constrained minimum-action paths). (ii) Forward simulation: for each branch path, we resampled 2,000 particles from the branch’s start-bin cells and evolved them by Euler–Maruyama integration of the learned field with the calibrated full-covariance noise (substeps ≤ 0.5 model time units). We sampled the simulated ensemble centroid at the observed time points, scored it, and normalized it by the constrained minimum action between the simulated path’s own endpoints. (iii) Random-partition centroids: for each branch, we centroided per-bin random cell sets of the same sizes (lineage structure destroyed, marginals preserved), scored them, and normalized them by their own-endpoint minima.

### 4.16 Morphogenetic-event and cell-cycle coupling tests

We assigned per-step action rates along the 179 worm paths to literature-curated windows (pharynx 330–430, muscle 430–480, enclosure 400–500, neural 420–560 min). For each window, we compared the observed fraction of total path action in the window to 1,000 circular time-shift permutations of the rate profile (pooled across paths). We adjusted p-values by Benjamini–Hochberg across the five windows. Division-rate coupling: per-path Spearman correlation between per-step action rate and the log fold-change in cell number of the path’s lineage between adjacent bins; significance by two-sided sign test over paths. Cell-cycle coupling: per-path Spearman correlation between action rate and a literature cell-cycle-length curve c(t) interpolated through (150, 30), (250, 35), (330, 55), (400, 75), (480, 90), (560, 100), (720, 110) min.

### 4.17 Perturbation analyses

In silico (worm): we ablated the rotational component of the drift, f_rot = f + ∇U, within 330–480 min by projecting per-step displacements onto −∇U. We recomputed observed-path actions and minimum-action paths for the 152 paths with window segments (Wilcoxon signed-rank test). Temporal rescaling stretched or compressed window segments by factors 0.5–2.0 before recomputing actions. Real perturbations: we subset the zebrafish perturbation atlas (Saunders et al. [29]; GEO GSE202639) to 240,000 cells spanning ctrl-inj and nine perturbation conditions (tbxta, tbx16, tbx16;msgn1, tbx16;tbx16l, cdx4, cdx4;cdx1a, noto, noto-mut, smo) at 18, 24, and 36 hpf (≥7 embryos per condition×timepoint). We trained a control field on ctrl-inj cells (PCA-50, same architecture). Per condition, the path was the chain of condition-pooled centroids across the three timepoints. We computed the action ratio under the control field with embryo-level bootstrap (100 replicates resampling embryos within condition×timepoint; 50 for the mesoderm-restricted variant). Point estimates used dual-initialization minimum-action optimization (300 iterations). Bootstrap replicates used a faster single-initialization variant (200 iterations), which slightly underestimates absolute ratios. We therefore assessed significance by a two-sample z-test comparing each perturbation’s bootstrap distribution to the control’s (standard errors from bootstrap interval widths), rather than by interval overlap with the point estimate.

External-anchor extension: we recomputed action ratios for all 14 conditions with full 18/24/36-hpf coverage (the nine above plus egr2b, epha4a, mafba, mafba-mut, wnt3a;wnt8) from a stratified 240,000-cell subsample of the all-genotype extraction (8,000 cells per condition×timepoint), with hours-consistent action units throughout field training, covariance calibration, and path scoring. Because the exact published subsample is unrecoverable, recomputed ratios shift in absolute scale (≈1.4× below published values) while preserving the published ranking of the nine overlapping perturbations (Spearman ρ = 0.78, p = 0.013, identical ranking under both optimization budgets below; cdx4;cdx1a is a persistent outlier: 3.84 published vs 1.9–2.3 recomputed). All anchor regressions therefore use recomputed ratios only. Severity metrics were derived from the published beta-binomial differential-abundance results: significant-test counts (n_sig), compositional total-variation distance (tv), Aitchison distance (aitch), and muscle-lineage displacement (muscle_d). The primary analysis correlated the ratio with each metric across the 14 conditions (Spearman and Pearson with bootstrap confidence intervals, Benjamini–Hochberg correction across the eight tests, leave-one-out cross-validated R²) under a 16-waypoint, 300-iteration full-metric optimization; the sensitivity arm used library defaults (40 waypoints, 500 iterations). The secondary analysis scored per-transition ratios (18→24 and 24→36 hpf; 33 transitions across 19 conditions, including five two-timepoint conditions) against end-timepoint n_sig with a condition-clustered bootstrap.

### 4.18 Temporal-resolution sensitivity

We re-scored worm paths on interpolated grids (0.25, 0.5, 1, 2 h; linear interpolation of centroid paths) and merged bins (1, 1.5 h). We re-scored zebrafish branches on the native stage grid, interpolated grids (0.5, 1, 2 h; full and post-epiboly windows), every-second-stage subsamples, and a uniform-1 h stage subset (7–12 hpf). All action integrals used per-step time increments.

### 4.19 Energetic translation of the action

We recovered raw UMI counts from the normalized checkpoint matrix. The stored matrix is log1p of size-factor-normalized counts, X = log1p(c·raw/SF), with the constant c = 6.048608 determined by matching per-cell sums to the authors’ UMI counts (median relative error 5×10⁻⁸); raw = expm1(X)·SF/c. For each of the 179 lineage anchors and each 30-min bin (≥10 cells; the same anchor–bin sets as the action analysis), we computed the per-gene raw-count centroid. We estimated the number of newly synthesized transcripts per cell per bin as Σ_g max(centroid_{b+1,g} − centroid_{b,g}, 0). This is a conservative lower bound: it counts only net increases and ignores mRNA degradation and dilution by cell division. A dilution-corrected upper variant used σ_g = max(x_{b+1} − x_b·2^(−d_b), 0), with d_b the embryo-wide doublings per bin from the cell-count curve below. Per-cell detected totals declined over the window (≈2,900 → ≈1,200 transcripts), consistent with decreasing cell size [20]. The embryo cell-count curve N(t) was piecewise log-linear through (100 min, 28 cells), (210 min, ≈190 cells), (700 min, ≈600-cell plateau), and (800 min, 558 cells) [20,28,30]. We converted transcripts to ATP equivalents at 2 (chain elongation only) to 48 (including de novo ribonucleotide biosynthesis) ATP equivalents per nucleotide [23], with a typical message length of 1.5 kb, and scaled by a 5– 20% detection efficiency [24]. We report both bounds and a central estimate (10% efficiency, geometric-mean per-nucleotide cost, dilution-corrected). We estimated total embryonic ATP turnover from oxygen consumption anchored to the mid-L1 basal rate (0.2 pmol O₂ min⁻¹ per animal [25]; embryonic mass-specific respiration is expected to be lower [31]) with P/O ≈ 2.5 (5 ATP per O₂). Time-resolved embryonic respiration is unavailable for *C. elegans*; existing embryo measurements are normalized within experiment [32]. For zebrafish, we unscaled the scaled matrix of the 3,000 highly variable genes with the stored per-gene means and standard deviations and exponentiated it to normalized (per-10⁴) counts. Lacking per-cell library sizes, zebrafish synthesis rates are relative, and we used them for profile and correlation analyses only. We digitized the zebrafish oxygen-consumption time course from Stackley et al. [26] (3–48 hpf). We recomputed per-step action rates as in 4.10. Action–synthesis associations used per-path Spearman coefficients (paths with ≥6 scored steps), exact sign tests, and pooled coefficients. The translation sensitivity used the lifetime protein:RNA cost ratio of 125–600× [23]. We computed mean profiles over bins with ≥20 contributing paths.

### 4.20 Field-speed metrics

For each observed path, minimum-action path, and straight-line route between the same endpoints, we evaluated the learned drift f at every waypoint. Per-path metrics: mean field magnitude |f|; mean traversal speed |Δx|/dt; the signed projection of f on the direction of motion, f·û; the fraction of steps with negative projection; the mean cosine between Δx and f; and a metric-consistent residual decomposition. For the decomposition, with W the Cholesky factor of the precision matrix (WᵀW = Σ⁻¹), we split the whitened residual W(ẋ − f) into components parallel and perpendicular to the whitened field Wf. We report them as fractions of the total quadratic form (which sums to the action integrand). Worm minimum-action paths used the settings of 4.6 (300 iterations, reproducing the reference minimum actions exactly). Zebrafish minimum-action paths were recomputed at a higher budget (1,200 iterations, best of straight-line and observed-path initializations). They descended a median 7% (max 27%; EVL/periderm branches) below the reference-budget values of 2.9, the same budget sensitivity noted in limitation (v). We compared observed and minimum-action-path metrics by paired Wilcoxon signed-rank tests per species (worm n = 179, zebrafish n = 25), with Benjamini–Hochberg adjustment across the seven metrics. Straight-line values are shown for reference.

### 4.21 Software and reproducibility

Analysis used Python 3.11 with scanpy 1.11, PyTorch 2.7.1, POT 0.9, scikit-learn, and SciPy. The LADD package and all analysis code are archived at Zenodo (https://doi.org/10.5281/zenodo.23540232). Random seeds were fixed throughout.

### 4.22 Statistical analysis

All tests were two-sided with a significance threshold of 0.05, and exact p values are reported throughout. Paired per-path comparisons used Wilcoxon signed-rank tests; sign consistency of within-lineage correlations used sign tests; monotonic associations used Spearman rank correlation. Enrichment of path action in morphogenetic windows and the external-anchor confirmatory endpoints were assessed by permutation tests (4.16; S9 Note). Where families of tests were computed, p values were adjusted by the Benjamini–Hochberg procedure (4.16, 4.20). Confidence intervals were estimated by bootstrap: a nested bootstrap over paths and model configurations for the headline action ratio (4.13), and an embryo-level bootstrap for the perturbation analyses (4.17). Analyses used SciPy as noted in 4.21.

### 4.23 Artificial-intelligence use disclosure

During the preparation of this work, the authors used an artificial intelligence (AI) research assistant (Biomni, Phylo [33]) to develop and execute analysis code under the authors’ direction, to generate figures, and to assist with language editing and restructuring of author-written text. All analyses were specified by the authors; all quantitative outputs were verified by the authors against the archived code and source-data tables (4.21), which reproduce every figure and number in the article. No primary data were generated by the AI tool, and no AI output was incorporated without author verification. The hypotheses, interpretations, and conclusions reported here are the authors’ own.

## Acknowledgments

Use of artificial-intelligence tools in the preparation of this work is disclosed in Methods 4.23.

## Author contributions

Qin Liu: Data curation, Formal analysis, Investigation, Validation, Visualization. Ying Zhou: Data curation, Formal analysis, Investigation, Validation, Visualization. Jielin Cao: Data curation, Formal analysis, Investigation, Validation, Visualization. Yiqiao Wang: Conceptualization, Methodology, Resources, Software, Project administration, Supervision, Writing – original draft, Writing – review & editing.

## Data and code availability

The *C. elegans* embryogenesis atlas is from Packer et al. [20] (GEO GSE126954); the zebrafish embryogenesis atlas from Farrell et al. [21] (GEO GSE106474; URD object from the Broad Single Cell Portal, SCP162); the zebrafish perturbation atlas from Saunders et al. [29] (GEO GSE202639). The LADD package, all analysis code, trained model checkpoints, and per-path source data tables are archived at Zenodo (https://doi.org/10.5281/zenodo.23540232; version-fixed snapshot) and are sufficient to reproduce every figure and table.

## Supporting information

S1 File. Supplementary notes S1–S12 and figures S1–S10.

S1 Fig. The learned developmental dynamics.

S2 Fig. Predicted futures respect the invariant lineage.

S3 Fig. LADD forecasts developmental dynamics out of sample.

S4 Fig. Forecasting robustness and diagnostics.

S5 Fig. Excess action tracks path geometry, not fate-map complexity.

S6 Fig. Exploratory localization of the excess action (post-hoc analyses).

S7 Fig. Forecasting benchmark.

S8 Fig. Additional action-analysis controls.

S9 Fig. Dimensionality trade-off and one-step prediction control.

S10 Fig. External severity anchor.

